# Linkage of nucleotide and functional diversity varies across gut bacteria

**DOI:** 10.1101/2025.06.06.658399

**Authors:** Veronika Dubinkina, Byron Smith, Chunyu Zhao, Cindy Pino, Katherine S. Pollard

## Abstract

Understanding the forces shaping genomic diversity within bacterial species is essential for interpreting microbiome evolution, ecology, and host associations. Here, we analyze over one hundred prevalent gut bacterial species using the Unified Human Gut Genome (UHGG) collection to characterize patterns of intra-specific genomic variability. Gene content divergence scales predictably with divergence in core genome single nucleotide polymorphisms (SNPs), though there is substantial variability in evolutionary dynamics across species. Overall, accessory genes exhibit consistently faster linkage decay compared to core SNPs, highlighting the fluidity of functional repertoires within species boundaries. This signal is strongest for mobile genetic elements, which show minimal linkage to core genome SNPs. Together, our findings reveal species-specific recombination regimes in the gut microbiome, underscoring the importance of accounting for horizontal gene transfer and genome plasticity in microbiome-wide association studies and evolutionary models.

## Introduction

The human gut microbiome is a highly diverse microbial community composed of Bacteria, Archaea, Eukaryotes, and viruses, which plays an essential role in health and disease^1–3^. Traditional, culture-based experimentation^4^, expanding reference databases^5–8^, and improved metagenomic tools^9–11^ have recently enabled strain-resolved analysis of this important community and have revealed enormous intraspecific diversity within and across human hosts^12–14^. In microbes, single nucleotide polymorphisms (SNPs) and larger differences in gene content can substantially affect physiology^15^, including driving differences in antimicrobial resistance^16–18^, pathogenicity^19^, and drug metabolism^20^. As a result, understanding the causes and consequences of this strain-level variation is critical in ecological and epidemiological studies of microbiomes.

One of the primary mechanisms affecting intraspecific diversity in bacteria is recombination, the movement of genetic material outside of transmission by linear descent^21,22^. Recombination can be conceptually divided into homologous and non-homologous processes. Homologous recombination involves the replacement of a genomic region with a highly similar sequence from a related strain or species^23,24^. This process, therefore, disrupts the strong linkage of alleles that occurs in linear descent and introduces variants into different genomic backgrounds^25^. Since homologous recombination requires highly similar sequences, it preferentially acts on the shared, core genome of a species^26–29^. Non-homologous recombination, on the other hand, can lead to transmission of accessory genes between strains^22^. This horizontal gene transfer (HGT) is often mediated by mobile genetic elements (MGEs) such as plasmids and viruses^30^. In the gut microbiome, where microbial species co-exist in dense and diverse communities, HGT may frequently and rapidly disseminate functional traits such as antibiotic resistance^31^, metabolic capabilities^32,33^, and virulence factors^34,35^. Homologous recombination and HGT rates can vary across species and genomic regions due to eco-evolutionary selective pressures and the molecular mechanisms facilitating them^26,36,37^. Together, these processes affect both the core and accessory genome, leading to rapid allele turnover^38,39^, phylogenetic signal erosion^40,41^, and the decoupling of gene content and SNPs^25,42^. While these processes have been studied in a few species of bacteria, specifically those with large numbers of cultured strains^28,43^, how they vary across species and shape populations of less-studied bacteria remains poorly understood.

Here, we explore patterns of intra-specific genomic variability in a large collection of reference genomes from bacterial species found in the human gut, the Unified Human Gut Genome collection (UHGG)^6^. We show that gene content divergence steadily increases with core genome divergence across more than a hundred of the best-represented gut bacterial species in the database. We estimate the decay of allelic linkage at increasing genomic distances both for SNP-SNP and accessory gene-SNP pairs, and use parameters of these curves to understand both recombination rates and population structure. We observe high variability of these parameters across species of gut bacteria. Despite the overall concordance between these two types of linkage-decay curves, we observed more rapid linkage decay for accessory genes, consistent with high rates of HGT. We confirm this intuition by showing that MGEs have almost no linkage to core genome SNPs in most species. Together, these results reveal that recombination and HGT shape species-specific patterns of genomic variation in the gut microbiome, with important implications for microbial evolution, ecology, and host-associated studies.

## Results

### Expanded reference genome collections capture strain-level variation in both the core and accessory genomes

Of the 4744 species represented in the UHGG, 121 had at least a hundred high-quality genomes (>95% completeness, <5% contamination) and were included in our analysis (Fig. 1D, Supplementary Table 1). For all species, we dereplicated closely related strains using the complement of the ANI (1 – ANI) – which we refer to here as the ANI-dissimilarity and use as a proxy for evolutionary divergence between pairs of genomes. After dereplication at a threshold of ANI-dissimilarity<0.001, we retained 42933 representative genomes, a median of 207 per species (interquartile range of 142 - 400) and about 15% of all UHGG genomes.

**Figure 1.**
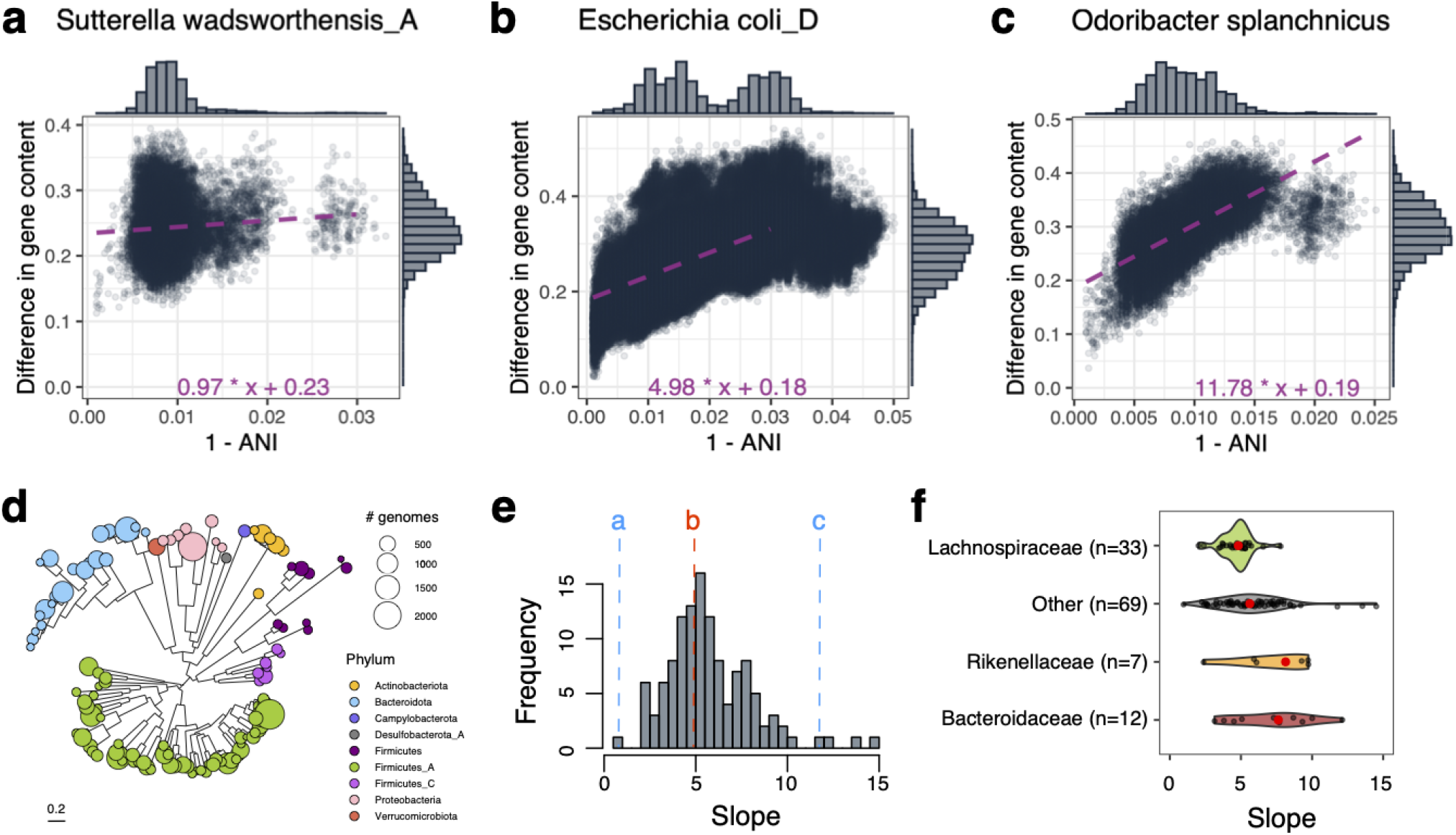
Evolution of gene repertoire of gut bacteria over time. **(a-c)**. Association between ANI-dissimilarity and gene content dissimilarity across pairs of genomes for three different species: **(a)** *Sutterella wadsworthensis* (n=205 genomes); **(b)** *Escherichia coli* (n=2259 genomes); **(c)** *Odoribacter splanchnicus* (n=333 genomes). The difference in gene content is calculated using binary distance for the presence/absence gene matrix. 1–ANI is calculated using skani (see Methods). We performed a linear fit (dashed line) on each plot using pairs within [0-0.03] of 1–ANI distance, as 0.03 is considered a species boundary. Marginal histograms indicate the distribution of points along each axis. **(d)** A phylogenetic tree of all species is included in our study. The color of the leaves represents phyla, and the size of the circles represents the number of genomes included in the analysis for a given species. **(e)** Distribution of slopes for linear fits across the entire collection of species (n=121). Dashed lines indicate values for the plots shown in (a-c). **(f)** Distributions of slopes for major taxonomic families in our dataset. Only families with significantly different slopes (p-value < 0.05) from the rest are shown.

The gene content of the UHGG collection is catalogued in the MIDAS reference database and has been previously clustered for each species at an 80% ANI clustering threshold [cite MIDAS]. Here we refer to these operational gene families as “genes” for simplicity. Using this resource, we estimated that reference genomes had a median of 2103 genes (IQR 1819 - 2495). Across all species, a median of 431 of these genes were found in >10% and <90% of genomes, a set which we refer to as accessory genes.

Analogous to ANI-dissimilarity, we calculated the pairwise gene content dissimilarity between genomes as the normalized hamming distance (“binary distance”). We found a high degree of gene content variation across species, with pairwise dissimilarities of 0.017 (median of per-species medians, IQR: 0.012 - 0.023). The size and diversity of this curated database of species, strain genomes, and accessory genes present an opportunity to study the evolutionary forces leading to strain-level functional variation in the human gut.

### Divergence in gene content increases with divergence in core-genome SNPs

We sought to understand how core genome and accessory genome coevolve. Comparing the accessory gene ANI-dissimilarity and SNP-dissimilarity (Fig. 1A–C), we observed a positive, linear relationship at <3% ANI-dissimilarity, although the slope and intercept of this linear relationship varied between species. For all species, the intercept was positive, indicating that even very closely related strains have significant gene content variation. The slope of this curve, while also universally positive (Supplementary Fig. 1), varied widely across species (Fig. 1E).

For some species (e.g., *Sutterella wadsworthensis_A*, Fig. 1A), the slope of this relationship was near 0, suggesting little concordance between SNP and gene content dissimilarity. For other species, like *Odoribacter splanchnicus* (Fig. 1C), the slope was much larger. *Escherichia coli*, as previously described^43^, is close to the median of this distribution (median slope = 5.30, *E. coli* slope = 4.98). Species in the family *Bacteroidaceae (*including the genus *Bacteroides)* and *Rikenellaceae* (including the genus *Alistipes)* have significantly higher genome plasticity, i.e., can rapidly gain and lose genes, while most of the bacteria belonging to the *Lachnospiraceae* have lower gene acquisition rates (Wilcoxon test p-value < 0.05; Fig. 1F, Supplementary Figure 1). However, even within bacterial families, there were considerable variations in the parameters of this relationship, signifying that ecological lifestyles may play a more prominent role in shaping the evolution of each species than taxonomy. There was a significant positive correlation between slope and genome size (Spearman correlation = 0.32, p-value < 0.001). This may reflect a shared common cause, as theoretical predictions suggest that for bacteria living in fluctuating environments, larger genomes and higher rates of HGT are both adaptive^44^.

### Core and accessory genomes and the different evolutionary forces shaping them

We next sought to determine whether SNPs in the core genome could be linked to the accessory genes in every species. To do this, we aligned individual strain genomes to a single representative genome for each species and identified core genome regions and biallelic SNPs (see Methods). We then calculated the pairwise SNP–SNP and accessory gene–SNP linkage (r^2^) as well as an estimate of the linear, genomic distance between loci. Next, we binned all pairwise LD estimates into specified distance ranges and calculated the 90th percentile of the pairs in each bin. We refer to this statistic as the LD90. Plotting the LD90 against genomic distance shows that linkage is reduced at longer distances (Fig. 2, Supplementary Figure 2).

**Figure 2.**
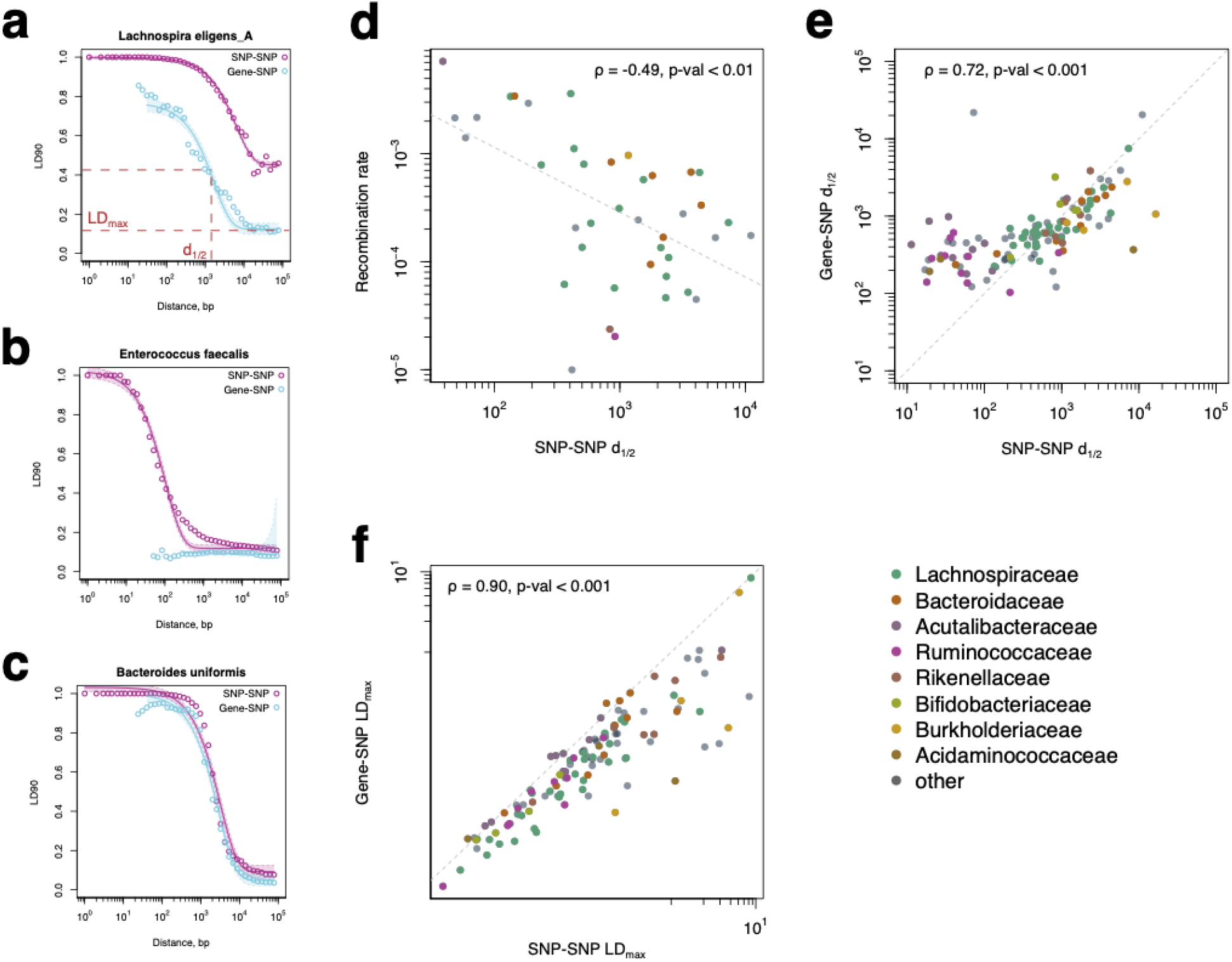
The linkage between SNP–SNP and accessory gene–SNP pairs. **(a-c)** Linkage decay curves for different gut bacterial species. The X-axis corresponds to the base pair distance on a chromosome between pairs of SNPs/accessory gene–SNPs (log-scaled and binned into 50 bins). The Y-axis shows each bin’s LD90 measure (90th quantile of correlation between all SNP–SNP or gene–SNP pairs). The line shows the exponential decay fit of each curve (see Methods). Shade indicates a 95% confidence interval for the fitted curve. This fit was used to infer two parameters: LD_max_, the value of LD90 at the maximal distance, and d1/2, the genomic distance at which linkage decays halfway. **(d)** Correlation between d_1/2_ for core SNPs linkage decay curves and recombination rates from ^39^. **(e)** Correlation between d_1/2_ for SNP– SNP and accessory gene–SNP linkage decay curves for different species. **(f)** Correlation between LD_max_ for SNP–SNP and accessory gene–SNP linkage decay curves for different species. Each point in (d) - (f) corresponds to a different species, the color of the points represents their taxonomic families, and the legend shows the Spearman correlation coefficient.

Comparing the linkage decay for accessory gene–SNP and SNP–SNP pairs and across species reveals several characteristic patterns: (i) Accessory gene–SNP linkage overall was significantly lower than SNP–SNP linkage within the core genome (Fig. 2A); (ii) Accessory gene–SNP linkage was much weaker than SNP–SNP linkage at shorter distances but remained consistent at longer distances (Fig. 2B); (iii) Accessory gene–SNP linkage was the same as SNP–SNP linkage in the core genome (Fig. 2C). The first two pattern types can be explained by additional disruption of linkage between accessory genes and SNPs as can be caused by frequent HGT. The latter signifies that the rate of HGT is either low or influenced by the species’ population structure.

We fit parameterized curves to these linkage-decay data (see Methods) and extracted two main statistics: d_1/2_ – the genomic distance at which correlation between pairs drops halfway– and LD_max_ – the baseline level of correlation between two pairs (Fig. 2A). For core genome SNP– SNP pairs, both parameters were driven mainly by homologous recombination: highly recombinogenic species are expected to have low d_1/2_ and low LD_max_^45^. In contrast, species with a substantial population structure disrupting rapid genetic transmission between lineages should have high LD_max_. HGT can also affect accessory gene–SNP pairs, disrupting the linkage between transferred genes and core genome SNPs. We confirmed this intuition by comparing our fitted parameters to recombination rates reported in a previous study of gut bacteria^39^. The SNP–SNP d_1/2_ had a significant negative correlation with a recombination rate (Spearman rho = –0.49, p-value < 0.01, Fig. 2D), while the accessory gene–SNP d_1/2_ was not significantly correlated with it (Spearman rho = –0.31, p-value > 0.05). Thus, by comparing the two curves, we can begin to disentangle these two processes shaping bacterial genomes.

Overall, there was remarkable variation in the d_1/2_ parameter for both SNP–SNP and accessory gene–SNP curves. Across gut bacteria, it ranged from 10bp to 10Kb, with no clear trend related to the phylogenetic relationships of the species (Fig. 2E). Interestingly, d_1/2_ for both SNP–SNP and accessory gene–SNP curves were strongly correlated (Spearman rho = 0.72, p-value < 0.001). There was a slight discrepancy between d_1/2_ at shorter scales, which could be attributed to the weaker statistical power for the accessory Gene–SNP curve, as we had fewer instances of gene-SNP pairs, making it more challenging to identify SNPs close to gene ends. Similarly, LD_max_ for both curves was also strongly correlated across species (Spearman rho = 0.90, p-value < 0.001, Fig. 2F). This supports the hypothesis that species population structure impacts the transmission of both core genome SNPs and accessory gene content.

### Patterns in core and accessory genome linkage across species

While the observed diversity in d_1/2_ and LD_max_ appears to indicate variation in both homologous recombination and HGT, we sought to better understand their relationships to a variety of other features of the underlying microbial species. We therefore calculated across all 121 species in our analysis all pairwise correlations among a set of continuous covariates, including parameters of the linkage decay curves, physiological traits, technical features of the available dataset, and genomic characteristics (Fig. 3A). As previously noted, accessory gene–SNP and SNP–SNP linkage decay curve parameters were strongly correlated (adjusted p-value < 0.01). LD curve parameters were also highly negatively correlated with the estimated average genome recombined fraction (i.e., positively correlated with the clonal fraction of the genome), which is consistent with our assumption that these statistics reflect recombination rates (adjusted p-value < 0.001). Notably, larger genomes (high number of genes) exhibited stronger linkage across both core and accessory genomes, and a smaller fraction of recombined genomes (adjusted p-value < 0.01). This is in line with a previous observation that smaller genomes tend to have higher gene turnover^43^. Higher GC content was associated with lower d_1/2_ and LDmax but a higher fraction of recombined genome, as shown before^46,47^. In addition, species represented by more strain genomes in the reference database and distributed across all continents exhibited lower d_1/2_ and LD_max_, as expected since those species likely have more opportunities for genetic exchange.

**Figure 3.**
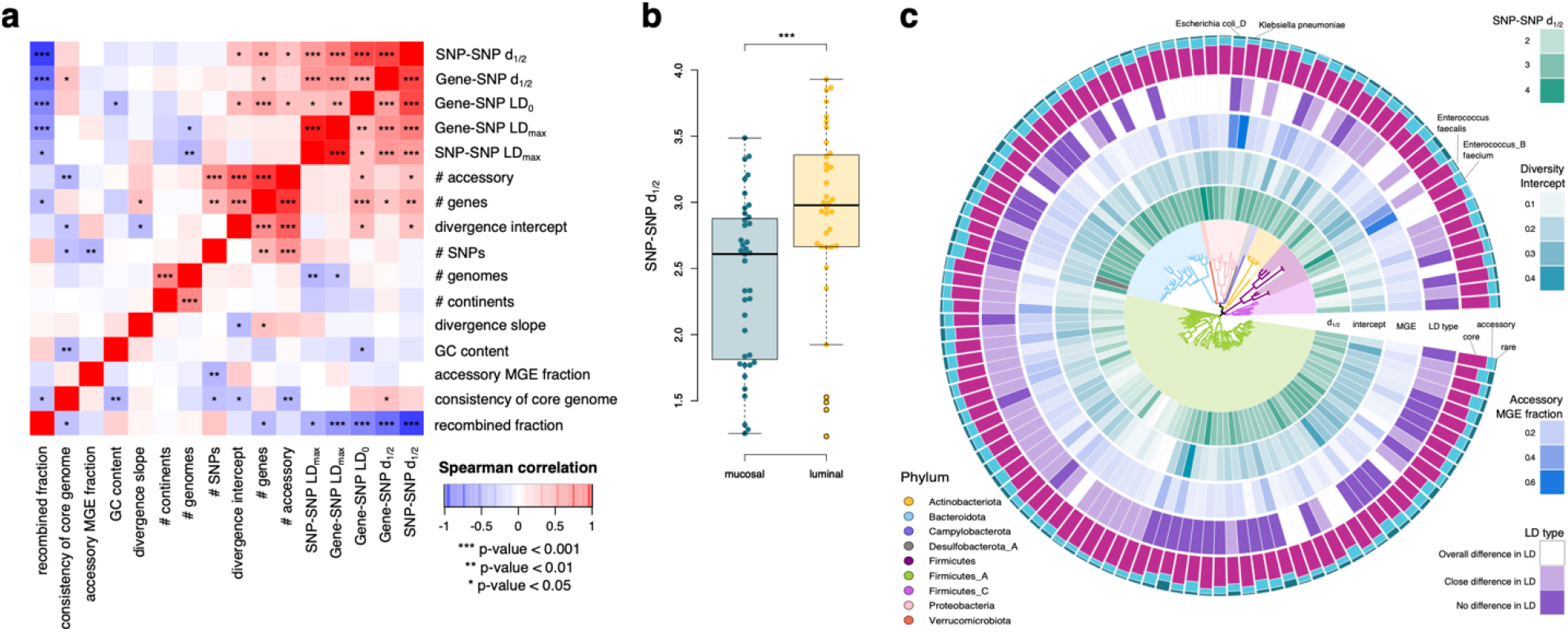
Global trends in parameters. **(a)** Heatmap of parameters’ correlations across the entire species set. We report five parameters from LD curves: SNP – SNP and accessory gene – SNP d_1/2_ and LDmax, described before, as well as gene – SNP LD_0_ – the value of LD90 at the zero distance; consistency of core genome is measured as the average pairwise agreement between gene order of all genomes to the reference; # accessory – the average number of accessory genes per genome of a given species, # genes – the average number of genes per genome of a given species; divergence intercept, slope – intercept and slope from the lines fitted in Fig 1; # SNPs – total number of *de novo* core genome SNPs in a given species; GC content – average GC content of genomes of the species; # genomes – number of genomes of a species included in the analysis; # continents – from how many different continents these genomes originated from according to UHGG metadata; accessory MGE fraction – average fraction of accessory genes that were annotated as a part of MGE across all genomes of the species; consistency of core genome – average relative number of breaks in gene order between the representative genome of each species and every other strain genome; recombined fraction – average fraction of recombined genome for a given species. We used Benjamini–Hochberg correction to adjust correlation test p-values. **(b)** Boxplot of d_1/2_ for SNP– SNP linkage decay curves for mucosal (n=40) and luminal (n=37) species. **(c)** Phylogenetic trends of some of these parameters. The species tree is colored by phylum. Heatmaps show the following parameters: d_1/2_ – SNP–SNP d_1/2_ (parameter representative of the LD curve parameters cluster on the heatmap in panel (a)); intercept – divergence intercept from fits in Fig 1 (representative of the second cluster in panel (a)); MGE – accessory MGE fraction; LD type – a categorical variable that indicates whether there is a significant difference in SNP-SNP accessory gene-SNP LD.

We also estimated the fraction of accessory genes that are annotated as MGE by geNomad^48^ and MobileElementFinder^49^ (see Methods). The MGE fraction was not significantly correlated with any parameters describing genomic diversity (i.e., LD curve parameters, Gene-ANI parameters, Fig. 3A). Additionally, we developed a measure of core genome gene order consistency to account for the fact that there might be some reshuffling across the bacterial chromosome between strain genomes of the same species (see Methods), which could significantly affect our estimation of distances between SNPs and genes. Note that the standard practice in population genetics is to assign SNP coordinates according to their coordinates in the reference genome. However, it remains to be determined to what extent reference genomes are “representative” of gene order in strains of that species overall. In our collection, most species had highly consistent core genome orders (mean consistency = 0.87 ± 0.03), and there was a slight positive correlation between this measure and higher clonal fraction of the genome.

We then looked for associations with bacterial lifestyle. We used phenotypic data available in the BacDive database^50^ as well as data from a previous study of luminal versus mucosal gut bacteria^51^ to annotate species in our collection (Supplementary Table 3) for binary lifestyle characteristics, and tested whether any of these features were associated with any of the observed statistics. We found no significant association of LD curve parameters to motile vs. nonmotile (n = 17 and 79 species, respectively), or sporulating vs. nonsporulating (14 and 82, respectively) species. However, there was a significant difference in LD for gram-negative vs. gram-positive (n = 39, n = 58) species (Wilcoxon rank-sum test, p-value < 0.01) and luminal vs. mucosal-associated (n = 37 and 40, respectively) species (Wilcoxon rank-sum test, p-value < 0.001, Fig. 3B). These two phenotypic variables are highly correlated (Chi-Square test p-value < 0.001), with a majority of luminal bacteria being gram-negative and mucosal being gram-positive. The higher d_1/2_ and lower recombination rates in gram-negative bacteria could be attributed to the fact that they have two membranes and a more complicated process of recombination/HGT^52^. In contrast, many gram-positive species are naturally competent^52,53^, and being in close proximity in mucosal biofilm gives them more opportunities to recombine^54^.

Next, we considered the taxonomic distribution of traits. Overall, there were no strong signals across the tree, with closely related species having very diverse parameters (e.g., *Enterococcus_B faecium* and *Enterococcus faecalis*). A notable exception was the strong separation between two major orders within *Firmicutes_A* phylum: *Lachnospirales* and *Oscillospirales*. While bacteria that belong to *Lachnospirales* had consistently lower d_1/2_ and almost no linkage even at close distances between accessory genes and core-genome SNPs, *Oscillospirales* had noticeably higher linkage and no difference between SNP–SNP and gene– SNP linkage curves (Wilcoxon rank-sum test, p-value < 0.001). However, even within these groups, there were examples of closely related organisms with very different parameter estimates. This result is corroborated by a previous study on recombination rates across bacteria, showing that while recombination rates generally appear to be a slowly evolving trait, they can evolve much faster in some genera^37^.

### Distinct heritability patterns across functional categories

We then looked into the variation in heritability of different functional categories in the accessory genome (i.e., the degree of linkage between core genome SNPs and genes). Certain types of genes that are subject to frequent HGT due to local selective pressures (e.g., prophages and insertion sequences, antibiotic resistance, or some capsid genes) are expected to be much less linked to the core genome population structure than others^55–57^. Indeed, for genes annotated as parts of MGEs (see Methods), the linkage decay curves were often flat (Fig. 4A, Supplementary Figure 2), with no association between these genes and core genome SNPs next to them.

**Figure 4.**
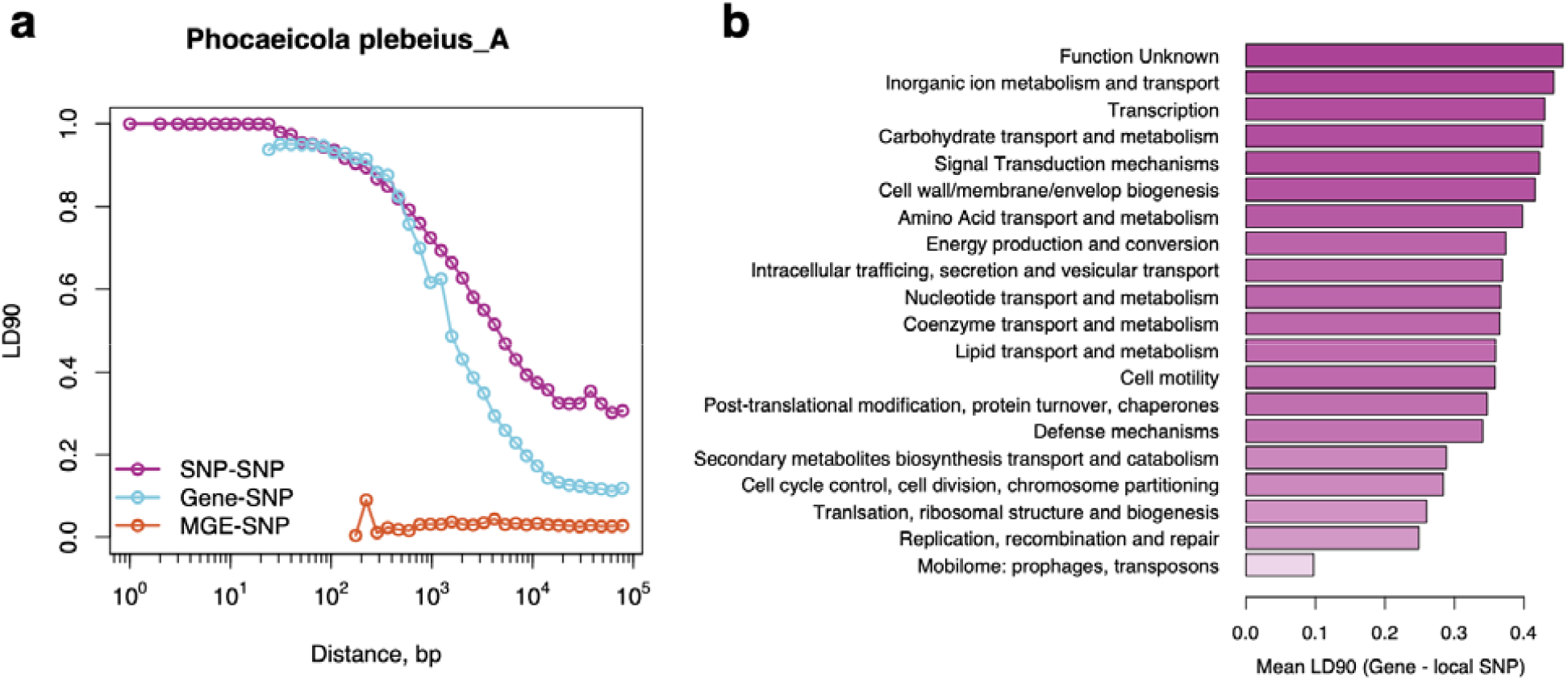
Variation in the heritability of genes with different functions. **(a)** Example of Linkage decay curves for mobile genetic elements genes (MGE) – SNP pairs. SNP–SNP and accessory gene–SNP curves are shown for comparison. The X-axis corresponds to the base pair distance on a chromosome between pairs of SNPs/accessory gene–SNP (log-scaled and binned into 50 bins). The Y-axis shows each bin’s LD90 measure (90th quantile of correlation between all SNP–SNP or gene–SNP pairs). **(b)** Barplot of per-category linkage. LD90 is calculated for all pairs of genes belonging to a given category with SNPs closer than SNP-SNP d_1/2_ (“local SNPs”) and averaged across all species.

We then stratified accessory genes by COG category and calculated LD90 for all genes within a given category and core genome SNPs that are closer than SNP–SNP d_1/2_ (designated “local” SNPs). In line with previous studies^6,58^, on average, metabolic functions were evolving more slowly (Fig. 4B), and thus were more linked to the core genome phylogeny than MGEs and some extracellular structure genes. However, there were noticeable differences in the heritability of functional categories across species (Supplementary Figure 3). For example, *Ruminococcus_E bromii* showed a high degree of linkage between MGEs and core genome SNPs, which might signify that these are “immobilized” MGEs (e.g., dead prophages). However, it is hard to make definite conclusions due to low gene counts in some COG categories and the few SNPs near them in some species. These findings suggest that the degree to which accessory genes are linked to the core genome varies not only by functional category but also by species-specific genomic and ecological contexts, reflecting diverse evolutionary constraints on gene mobility within the gut microbiome.

## Discussion

The interplay between the evolution of the core and accessory parts of bacterial pangenomes remains a central topic in microbial population genomics^35,59,60^. While it has been studied to some extent how homologous recombination shapes the core genomes of different species^27,29,37,39^, it is still unclear how accessory genes fit in this picture. With the recent expansion of bacterial genome collections^5–7,50,61^, we can address these questions in more detail beyond the scope of a few well-studied bacteria, such as *E. coli*^43^ *and Streptococcus pneumoniae*^62^. In this study, we comprehensively analyzed core and accessory intra-species diversity for the 121 most prevalent human gut bacteria. Our analyses reveal a consistent trend of increasing gene content divergence with core genome divergence across a broad range of prevalent gut bacterial species, suggesting a tight coupling between vertical inheritance and gene gain/loss, potentially reflecting long-term evolutionary trajectories shaped by selection and ecological niche differentiation.

By estimating the decay of allelic linkage with genomic distance, we probed recombination dynamics within species using both SNP-SNP and accessory gene-SNP linkage profiles. These curves capture how rapidly linkage disequilibrium breaks down—a function of both recombination rate and population structure. We observed considerable heterogeneity across species in the shape and steepness of these decay curves, underscoring the diverse evolutionary histories and recombination regimes among gut microbes.

One observation of our study is that we found a high degree of concordance between linkage decays for core SNP–SNP and those for accessory gene – core SNP pairs (Spearman rho = 0.72, p-value < 0.001). This might signify that, in gut bacteria, the evolution of core and accessory genomes is affected by the same types of selective pressures and molecular mechanisms (i.e., recombination boundaries are similar to HGT boundaries). When we compared genomic distances at which the correlation between SNP–SNP and accessory gene– SNP pairs drops halfway, we noted that it can vary a few orders of magnitude between just a hundred base pairs and ten Kbp. Species with a high degree of linkage tend to have larger and more consistent genomes, lower recombination rates, and lower GC content. Most of them tend to show geographic restriction, are gram-negative, and enriched in luminal fraction, which might explain why it is harder for them to exchange genetic material. Meanwhile, only a few species in our collection exhibit a strong population structure that manifests as a high degree of linkage at long genomic distances: *Phocaeicola dorei, Enterococcus_B faecium, Parasutterella excrementihominis*, and *Ruminococcus_E bromii*.

Notably, while the decay profiles of SNP-SNP and gene-SNP linkages were broadly concordant, accessory gene linkages decayed more rapidly in nearly all cases. This suggests that HGT, particularly of MGEs, plays a dominant role in shaping the accessory genome. Our finding that MGEs are almost entirely unlinked to core genome SNPs further supports this, indicating their movement is largely independent of vertical descent or homologous recombination involving the core genome.

We also discovered that for about a third of the species in our collection, there is a significant difference in short-distance linkage between SNP–SNP and accessory gene–SNP pairs. While it could be a consequence of low number statistics (only a few genes have nearby SNPs), another intriguing possibility is that the accessory genomes of these species are primarily shaped by MGEs. Indeed, as expected, genes that are annotated as parts of MGE have almost no linkage to core genome SNPs. However, we did not find any correlation between the fraction of MGEs in the accessory genome and linkage patterns across our species collection. This might be due to the inconsistency of MGE annotations across different species.

These observations have several important implications. First, they reinforce the idea that accessory genes—often involved in functions such as antibiotic resistance, phage defense, or metabolic flexibility—are frequently decoupled from core genome evolution. However, as evident in our analysis, this is true only for a subset of gut bacteria. This has consequences for how we interpret gene-based associations in microbiome-wide studies: the presence of a particular gene may not imply a recent common ancestor between strains, and assumptions of clonal inheritance may be misleading when accessory functions are mobile.

Second, our approach illustrates how linkage decay patterns can serve as a scalable, genome-wide readout of recombination and population structure, even in species with limited cultivation or with high levels of within-host diversity. The distinction between SNP-SNP and gene-SNP linkage decay provides a practical way to separate homologous recombination from gene flow via HGT, helping disentangle evolutionary processes operating on different timescales and genomic units.

Finally, the striking variability in linkage decay and gene content dynamics across species phylogeny highlights the importance of species-specific analyses in microbial population genetics. Unlike model organisms or well-studied pathogens, many gut commensals lack standardized population genetic models. Our findings argue against one-size-fits-all assumptions and call for more nuanced frameworks integrating selection, recombination, and gene mobility across taxa. Variations in the recombination statistics estimated in our study could reflect variable selection pressures across ecological niches, differences in fundamental genomic biology, or, in some cases, bias introduced by variable dataset size and data quality. Unfortunately, our understanding of gut bacterial ecology and phenotypes remains very limited, hindering efforts to disentangle these factors.

Looking forward, several open questions remain. How do host factors such as diet, immune state, or geography shape recombination landscapes across individuals? Can linkage decay curves be extended to metagenomic samples to infer recombination in situ? What evolutionary forces maintain high rates of gene mobility in some species but not others? By combining large-scale genomic data with population genetic modeling, our work provides a foundation for answering these questions and for incorporating recombination-aware methods into microbiome analysis pipelines. As the field moves toward higher resolution, strain-level microbiome profiling, accounting for gene mobility and recombination will be essential for robust inference of microbial function, evolution, and association with host phenotypes.

## Methods

### Dataset construction

We retrieved the UHGG v2 database^6^, consisting of 289232 genomes and 4744 species. Since the majority of the genomes in UHGG are MAGs that could contain sequencing errors and assembly artifacts, we focused our analysis on species with at least 100 high-quality genomes (>95% completeness and <5% contamination), leaving us with 74614 genomes across 158 species. The genome with the best quality is typically assigned as a representative one for each species. We used skani v0.2.2 with default parameters^63^ to estimate all-to-all ANI between strain genomes of each species and left only those with ANI>99.9 to any other genomes. This left us with 121 species and 42933 genomes (∼15% of the UHGG database), the final set of genomes used in this study (Supplementary Table 1). We used the phylogenetic tree provided by the authors of UHGG for all phylogenetic analyses.

### Pangenome analysis

We used gene clusters as defined in the MIDAS v3 database^11^. Briefly, for each reference genome assigned to a given species in the UHGG v2 genome collection, genes were predicted by Prokka v1.14.6^64^. All gene sequences less than 200bp or with ambiguous bases were filtered out. The remaining sequences were dereplicated using VSEARCH v2.23.0^65^, using a 99% ANI cut-off, and the longest sequence was assigned as the representative for each cluster. In addition, we removed fragmented genes by running CD-HIT v4.8.1^66^ (using options ‘-c 1 -aS 0.9 -G 0 -g 1 -AS 180’) and merging all clusters with shorter representative sequences having the perfect identity of ≥ 90% of length to another longer cluster representative (same applies to gene sequences predicted on the opposite strand^67^). We then clustered representative sequences into OGFs using VSEARCH and an 80% gene similarity cut-off. We used binary presence/absence for genes (ignoring the multi-copy genes, as they constitute only ∼5% of the genes).

We used the presence/absence of these gene clusters within every genome to calculate pairwise gene differences for every species (Fig. 1). For subsequent analyses, we defined “core” genes as genes present in >90% of the strains, “rare” as present in <10% of the strains, and the rest were assigned as “accessory”. We estimated the consistency of the core genome by calculating the average relative number of breaks in gene order between the representative genome of each species and every other strain genome.

We annotated gene sequences using EggNOG mapper v2.1.12^68^. We run geNomad v1.7.4^48^ and MobileElementFinder v1.1.2^49^ directly on every reference genome for a species to identify mobile genetic elements (phages and plasmids). We used the result of this analysis to calculate the average fraction of accessory MGE genes per species by checking which accessory genes were predicted to be part of MGE in every given genome.

### SNP profiling

For each species, we mapped all genomes against the species’ representative genomes (in UHGG, this is the genome with the highest assembly quality, often an isolate) using ‘nucmer’ from MUMmer v4.0.0rc1^69^. We used ‘show-snps -ClrT’ and ‘show-coords -rgT’ to identify SNPs and extract their coordinates. Only bi-allelic SNPs with at least 5% minor allele frequency were included in the catalog, and we required that they be covered in at least 90% of the genomes (to include only core-genome positions). In the final SNP catalog for each species, we marked alleles as “major,” “minor,” and “NA” (position not covered) and identified them by their coordinates in the representative genome.

### Linkage decay estimation

To estimate the SNP–SNP linkage decay curve, we defined 50 bins in the interval [0-100Kb] using logarithmic scaling ([0,1), [1,2), [2,3), … [48077, 61370), [61370, 78339), [78339, 100000)). The distance between a pair of SNPs was calculated using their coordinates in the reference genome. We then calculated the 90th quantile of pairwise SNP correlations within each bin (SNP pairs from different contigs were excluded in this calculation). To estimate the accessory gene–SNP linkage decay curve, we used the same 50 bins. To calculate the distance between a given gene and SNP, we extracted all minimal genomic distances between gene ends and SNP positions when detected in the same strain genome within the same contig and calculated the average distance across all such genomes. Note that this measure of genomic distance is slightly more robust than the distance defined on the reference genome for a few reasons: only a subset of accessory genes is present in any given reference genome; reference genome coordinates for SNPs and genes might not be representative due to reshuffling of gene order between genomes (as estimated by the consistency of core genes order). We used the following formula to approximate linkage decay curves:

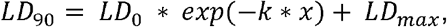

from this fit, we extracted two main parameters: LD_max_ and 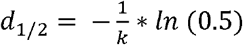.

### Recombined fraction estimation

We used a procedure similar to the recent study^39^ to estimate the fraction of the recombined genome. We used our de novo core genome SNP data to estimate SNP density across the genome, split into 500bp windows. We then modeled the distribution of SNP density as a Poisson + Negative Binomial Mixture Model and used these probabilities to classify all windows into “clonal” and “recombined” states. We used these labels to initialize a two-state HMM model that assigns final labels for “clonal” and “recombined” parts of the genome. Throughout the study, the fraction of the recombined genome for a given species was calculated as an average fraction of “recombined” windows across all pairs of strains with the reference genome.

## Statistical data analysis

We used R 4.2.1 for statistical analysis. We used *nlsLM* from the package *minpack*.*lm* for linkage decay curve fitting; *cor*.*test(method=“spearman”)* from the *stats* package to calculate correlation and p-value for parameter associations; Wilcoxon test for all pairwise distributions comparisons, and Benjamini-Hochberg adjustment for p-value correction.

## Data and code availability

All the genomes included in the analysis are listed in Supplementary Table 1 and can be retrieved from the UHGG database. The code used to reproduce the data analysis and figures is available here: https://github.com/Zireae1/snp2genes. Any additional information required to reanalyze the data reported in this paper is available from the lead contact upon request.

## Acknowledgments

This work was supported by NHLBI (grant #R01HL160862 to KSP), the San Simeon fund, Chan Zuckerberg Biohub SF, Gladstone Institutes, and an NSF predoctoral fellowship to CP. This research was supported in part by grant NSF PHY-2309135 and the Gordon and Betty Moore Foundation Grant No. 2919.02 to the Kavli Institute for Theoretical Physics (KITP).

## Authors contributions

VD: Conceptualization, Methodology, Software, Formal Analysis, Investigation, Writing – Original Draft, Writing – Review & Editing, Visualization

BJS: Conceptualization, Methodology, Software, Formal Analysis, Investigation, Writing – Original Draft, Writing – Review & Editing, Visualization

CZ: Methodology, Software, Writing – Original Draft, Writing – Review & Editing CP: Writing – Review & Editing

KP: Conceptualization, Methodology, Investigation, Resources, Writing – Original Draft, Writing

– Review & Editing, Supervision, Funding Acquisition

## Notes

### Competing Interest Statement

The authors have declared no competing interest.

