## Supplementary figures 1-3 for "Linkage of nucleotide and functional diversity varies across gut bacteria"

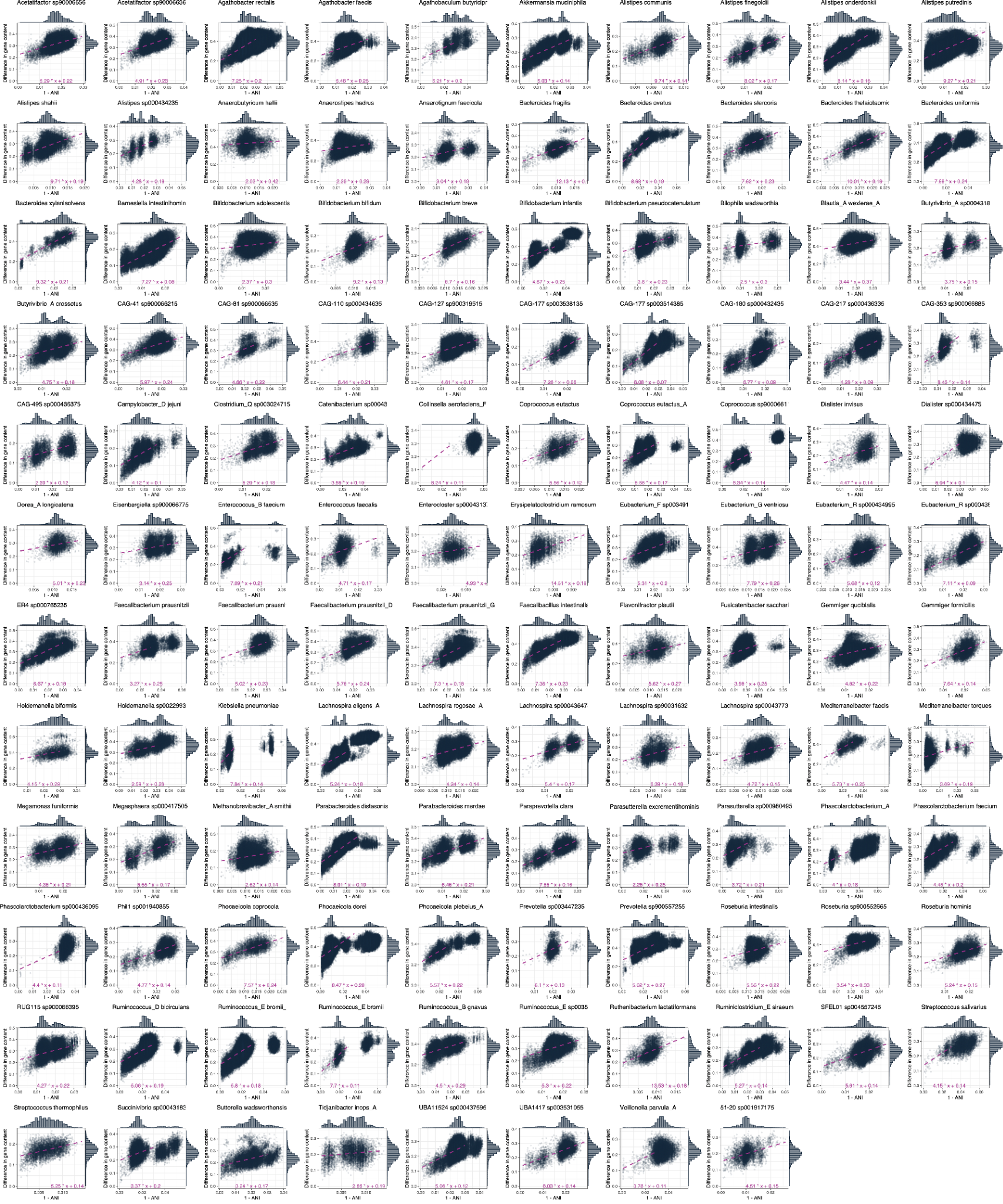

**Supplementary Figure 1.** **Evolution of gene repertoire of gut bacteria over time.** Association between gene content divergence and genome nucleotide divergence across pairs of genomes for all species in our analysis.

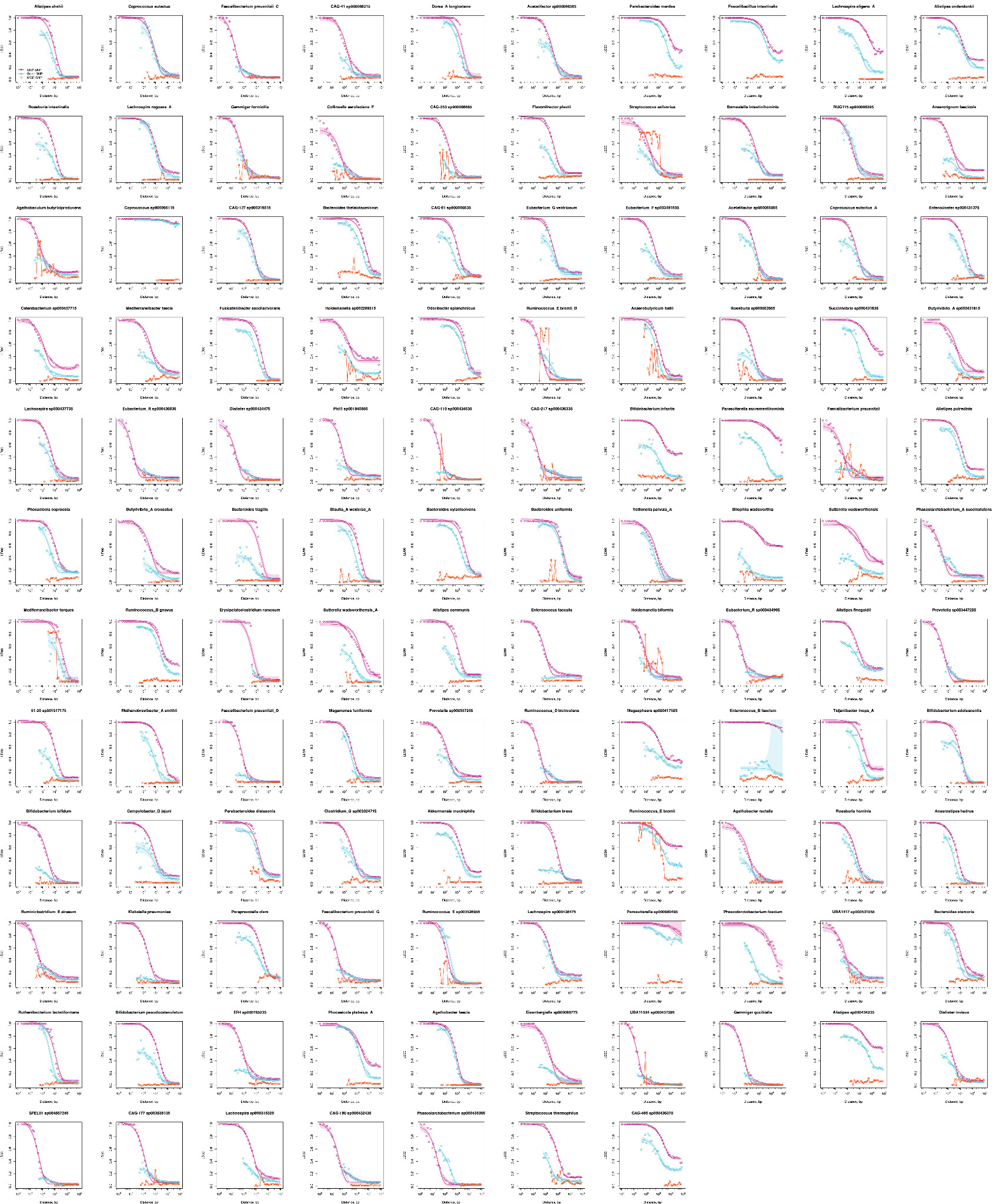

**Supplementary Figure 2. LD decay curves for all species for SNP–SNP (purple), gene–SNP (blue), and MGE–SNP (orange) pairs.** The shaded part shows a 95% confidence interval for the fitted curves for SNP–SNP and gene–SNP decays.

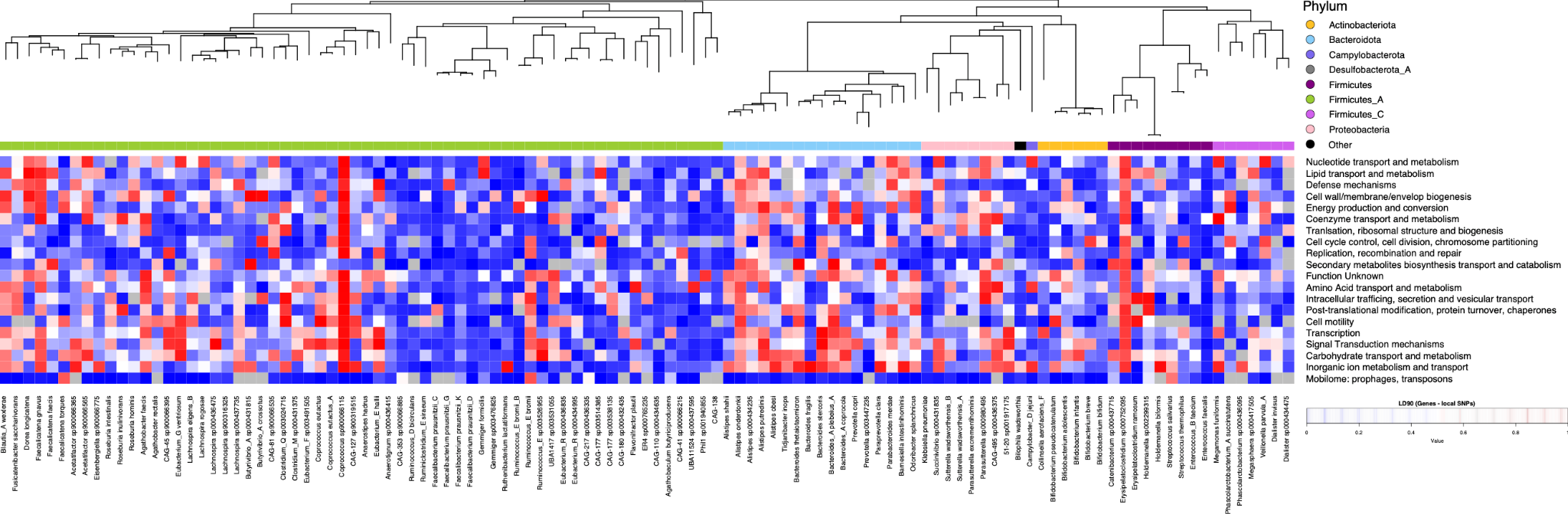

**Supplementary Figure 3. LD90 between accessory genes in 20 major COG categories and “local” core genome SNPs across species.** Grey color shows cases where no accessory genes are assigned to a given category.
